# KRAB Zinc Finger Proteins coordinate across evolutionary time scales to battle retroelements

**DOI:** 10.1101/429563

**Authors:** Jason D Fernandes, Maximilian Haeussler, Joel Armstrong, Kristof Tigyi, Joshua Gu, Natalie Filippi, Jessica Pierce, Tiffany Thisner, Paola Angulo, Sol Katzman, Benedict Paten, David Haussler, Sofie R Salama

## Abstract

KRAB Zinc Finger Proteins (KZNFs) are the largest and fastest evolving family of human transcription factors^1,2^. The evolution of this protein family is closely linked to the tempo of retrotransposable element (RTE) invasions, with specific KZNF family members demonstrated to transcriptionally repress specific families of RTEs^3,4^. The competing selective pressures between RTEs and the KZNFs results in evolutionary arms races whereby KZNFs evolve to recognize RTEs, while RTEs evolve to escape KZNF recognition^5^. Evolutionary analyses of the primate-specific RTE family L1PA and two of its KZNF binders, ZNF93 and ZNF649, reveal specific nucleotide and amino changes consistent with an arms race scenario. Our results suggest a model whereby ZNF649 and ZNF93 worked together to target independent motifs within the L1PA RTE lineage. L1PA elements eventually escaped the concerted action of this KZNF “team” over ∼30 million years through two distinct mechanisms: a slow accumulation of point mutations in the ZNF649 binding site and a rapid, massive deletion of the entire ZNF93 binding site.

---

KZNFs repress RTEs by recruiting the co-factor Kap1 (Trim28) which then recruits a variety of repressive factors that establish heterochromatin^1,6^. We reasoned that KZNFs expressed highly in the pluripotent stem cell (PSC) state were likely to be involved in arms race scenarios as the pluripotent state is a high stakes evolutionary battleground since RTEs that successfully retrotranspose in this state are inherited by all daughter cells including the germ line^7^. We further narrowed our investigation to the L1PA RTE lineage, which contains the only active, autonomous RTE family (L1HS elements) in humans, and is therefore likely to experience high evolutionary pressure for repression^8^. Recent ChIP-SEQ studies have revealed many KZNFs that bind L1PA families (Extended Data Fig 1), although these studies were performed in an artificial overexpression context in 293T cells^4,9^. In order to analyze which of these binders might be important for repression in the PSC context, we mapped KZNF ChIP-SEQ data to consensus repeat elements via the UCSC Repeat Browser (*MH, in preparation*), and then correlated Repeat Browser “meta-peaks” with Kap1 ChIP-SEQ “meta-peaks” from PSCs (Fig 1A). This analysis identified two KZNFs, ZNF649 and ZNF93 which are highly expressed in the PSC state (Extended Data Fig 1), as responsible for the majority of Kap1 recruit on L1PA elements in the human PSC context. We previously identified ZNF93 as an important repressor of L1PA elements in hPSCs, and traced its binding site to a 129-bp region that was deleted in the youngest (L1PA2, L1HS) L1PA families^5^. Interestingly, ZNF649 recognition of L1PA elements is strongest on L1PA6-L1PA4 elements, appears to weaken in younger elements (L1PA3-L1PA2), and is unable to bind L1HS elements (Fig 1A). Additionally, the correlation between Kap1 binding and ZNF649 and ZNF93 binding also varies across each L1PA family, suggesting that ZNF93 is most effective on L1PA4 and L1PA3, while ZNF649 was most active on older L1PA5 elements but slowly lost its efficacy (Fig 1A). However, the DNA changes that allowed L1PA escape from ZNF649 require more complex analysis than the ZNF93 case.

**Figure 1:**
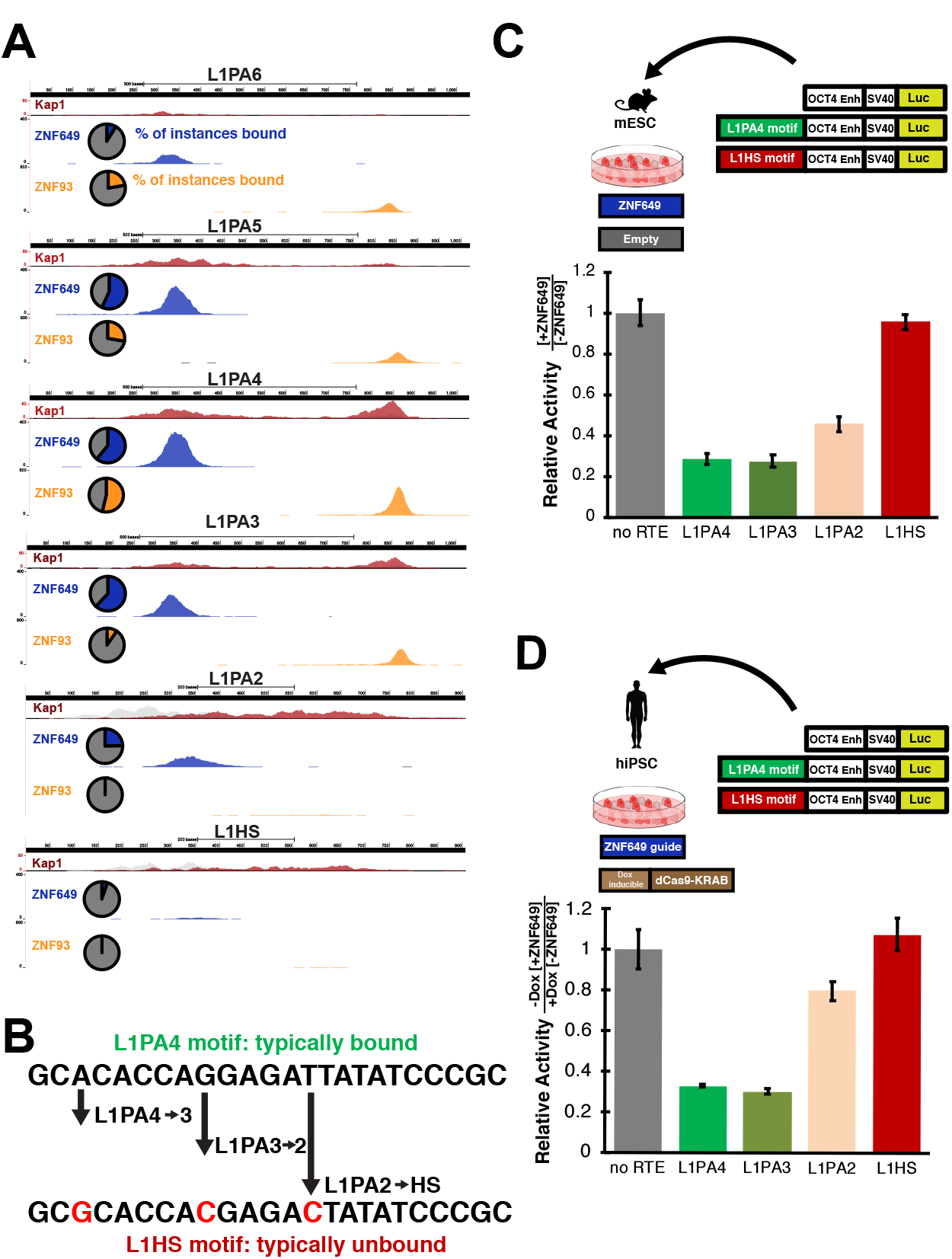
ZNF649 recognizes a sequence-specific motif in L1PA elements. A) Repeat Browser Analysis of ZNF649 (blue), ZNF93 (gold), Kap1 (red/grey; independent replicates) on L1PA elements. Pie charts show the percentages of each L1PA family that are independently bound by ZNF649 (blue slices) and ZNF93 (yellow slices). B) Discriminative analysis of the sequence under the ZNF649 binding site reveals a recognition motif that acquires 3 sequential point mutations in younger L1PA families. C) Reporter assay in mouse embryonic stem cells (mESC) in which a reporter containing the ZNF649 binding site in each of the L1PA sequences are tested in the presence or absence of ectopic ZNF649. Bars represent the relative activity of the reporter in the presence of ZNF649 normalized compared to an empty vector. D) Analogous experiment in CRISPRi hiPSCs expressing guides targeting ZNF649 and inducible dCas9-KRAB. Induction with dox depletes endogenous ZNF649. Data represents reporter activity in the knockdown condition normalized by the uninduced endogenous condition (all activities normalized to no RTE control). All error bars represent standard deviations of four biological replicates.

In order to determine how L1HS escaped ZNF649 binding, we compared the sequences^10^ (centered around our Repeat Browser meta-summits) of L1PA elements with ChIP-SEQ summits versus that of L1PA elements without summits. This analysis revealed two adjacent motifs that together form one long putative ZNF649 recognition sequence (Fig 1B). To test if this sequence was sufficient for ZNF649 binding, we transfected a reporter plasmid with the ZNF649 binding site cloned upstream of a luciferase gene in the presence or absence of human ZNF649 in mouse embryonic stem cells (mESCs) which are free of other primate-specific repressive elements^5^. ZNF649 specifically repressed this sequence, validating it as a bona fide recognition motif (Fig 1B). We then examined the evolution of this motif in the L1PA families which appear to escape ZNF649 recognition. The ZNF649 recognition sequence accumulated three mutations that flourished in younger families over the last 18 million years (Fig 1B). We tested the effect of each of these mutations (representing L1PA3, L1PA2 and L1HS-like states) and observed complete loss of repression in the L1HS-like state and an intermediate phenotype in the L1PA2-like state. To confirm that our reporter system accurately recapitulated ZNF649 action within a human PSC, we performed CRISPRi knockdowns of ZNF649 in human iPSCs^11^ and repeated our reporter assay. These experiments matched our mESC results, with the reporter repressed by endogenous ZNF649 but expressed in the knockdown context (Fig 1C).

Interestingly, the ZNF649 recognition motif contains a TATA sequence^12^, an important recognition motif for the transcription factor TBP. Furthermore, this TATA sequence arose in a subset of the L1PA6 lineage through a dramatic rearrangement of the L1PA 5’ UTR that also deleted an old TATA sequence present in ancient elements (Fig 2A). While only a minority of L1PA6 elements contained this new arrangement of the 5’UTR, almost all instances of younger L1PA elements are configured in this manner suggesting a comparative fitness advantage (Fig 2A). Furthermore, mutation of the TATA sequence results in a complete loss of ZNF649-mediated repression (Fig 2B), demonstrating a potential mutational route for L1PA escape from ZNF649. However, mutations in this site are rare indicating that L1PA elements have a competing selective pressure (presumably to maintain transcription factor binding to initiate transcription) to avoid this route.

**Figure 2:**
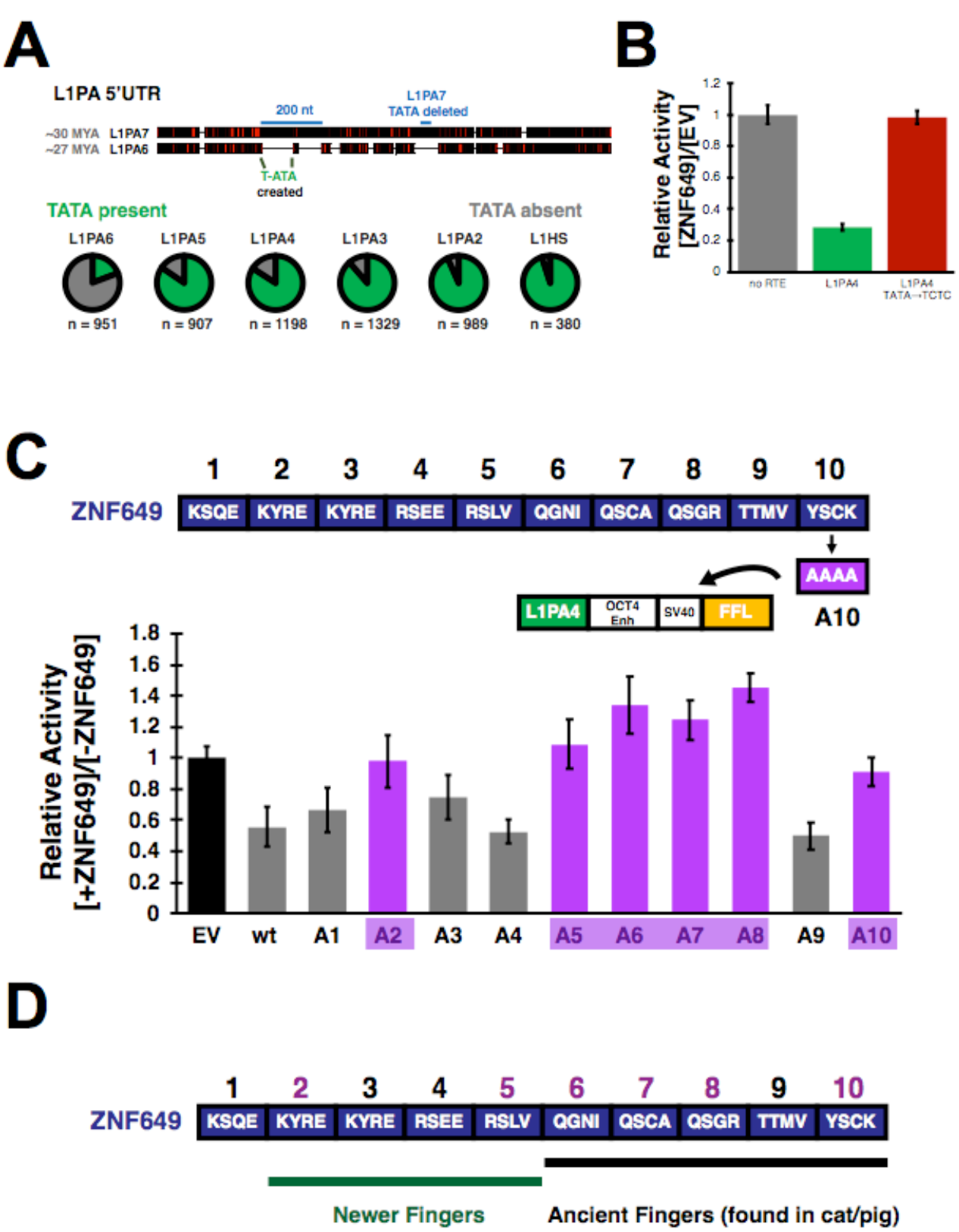
Evolution of L1PA families and ZNF649. A) Consensus L1PA7 sequence aligned to consensus L1PA6 sequence. Black coloring indicates conservation, red indicates variation. *(Below)* Pie charts representing the number of instances of each L1PA family that have a perfectly conserved “TATA” sequence (green slice) in the new 5’ configuration. B) mESC reporter assay measuring ZNF649’s ability to repress when the highly conserved TATA sequence is mutated. C) Testing of finger mutations in the mESC reporter assay. Shown is the relative activity of each single finger mutant (A1-A10) compared to wild type (wt). Purple coloring indicates mutants that show no repression. D) Cartoon representation of ancient and modern fingers of ZNF649 overlayed with reporter data from C).

In order to understand how ZNF649 evolved to repress these elements, we synthesized 10 ZNF649 mutants (one for each ZNF domain) with each mutant designed to ablate the binding activity of a single finger. Canonically each ZNF within a KNZF recognizes three nucleotides of double stranded DNA via four specific DNA contact residues, typically (though not always) amino acids –1,2,3 and 6 relative to the ZNF helix^13,14^ (Extended Data Fig 2); therefore, we mutated all four canonical DNA contact residues to alanine in each construct and tested each mutant’s ability to repress the L1PA4 luciferase reporter. Mutations to fingers 2,5,6,7 and 8 led to a loss of repression, indicating their importance in L1PA recognition. (Fig 2C). We then traced the evolutionary history of ZNF649, which is found across Eutheria, making it over 100 million years old. Determining true orthologs of KZNFs is challenging given their rapid evolutionary rates, high sequence similarity, propensity for duplication, and potential for gene conversion. By focusing on individual ZNF domains we reconstructed ancestral states for ZNF649 in the primate lineage *(Armstrong et al., in preparation)*. L1PA6 elements with the modern 5’UTR configuration are found only in Old World and not New World monkeys, meaning that ZNF649 must have evolved to battle these RTEs within the last 30 million years. Fingers 2 and 5, identified by our mutational analyses as being critical for L1PA repression, occur in a part of the gene that appears to have evolved more recently *(Armstrong et al, in preparation)*. Fingers 6, 7, and 8 which are also critical for binding, have more ancient roots, clearly matching fingers in distantly related species such as pigs and cats (Fig 2D, Extended Data 3). These fingers are bioinformatically predicted to bind the TATA sequence^15^ which may suggest an ancient gene regulatory role for ZNF649 at a TATA sequence; ZNF649 may have then been repurposed to target the TATA sequence of L1PA6 and younger elements upon RTE invasion, which resulted in the acquisition of fingers 2 and 5.

Together these data suggest a model whereby L1PA elements rearranged their 5’UTRs ∼30 million years ago – perhaps to escape repression from an unknown ancient KZNF. ZNF649 quickly adapted to bind L1PA elements at the new TATA sequence by gaining new fingers, followed by rearrangements in ZNF93 that allowed it to repress L1PA4 elements. When L1PA3 elements responded by deleting the entire ZNF93 binding site, ZNF649 was left to battle L1PA elements on its own – a battle that it then lost as mutations accumulated to generate the active L1HS state (Fig 3).

**Figure 3:**
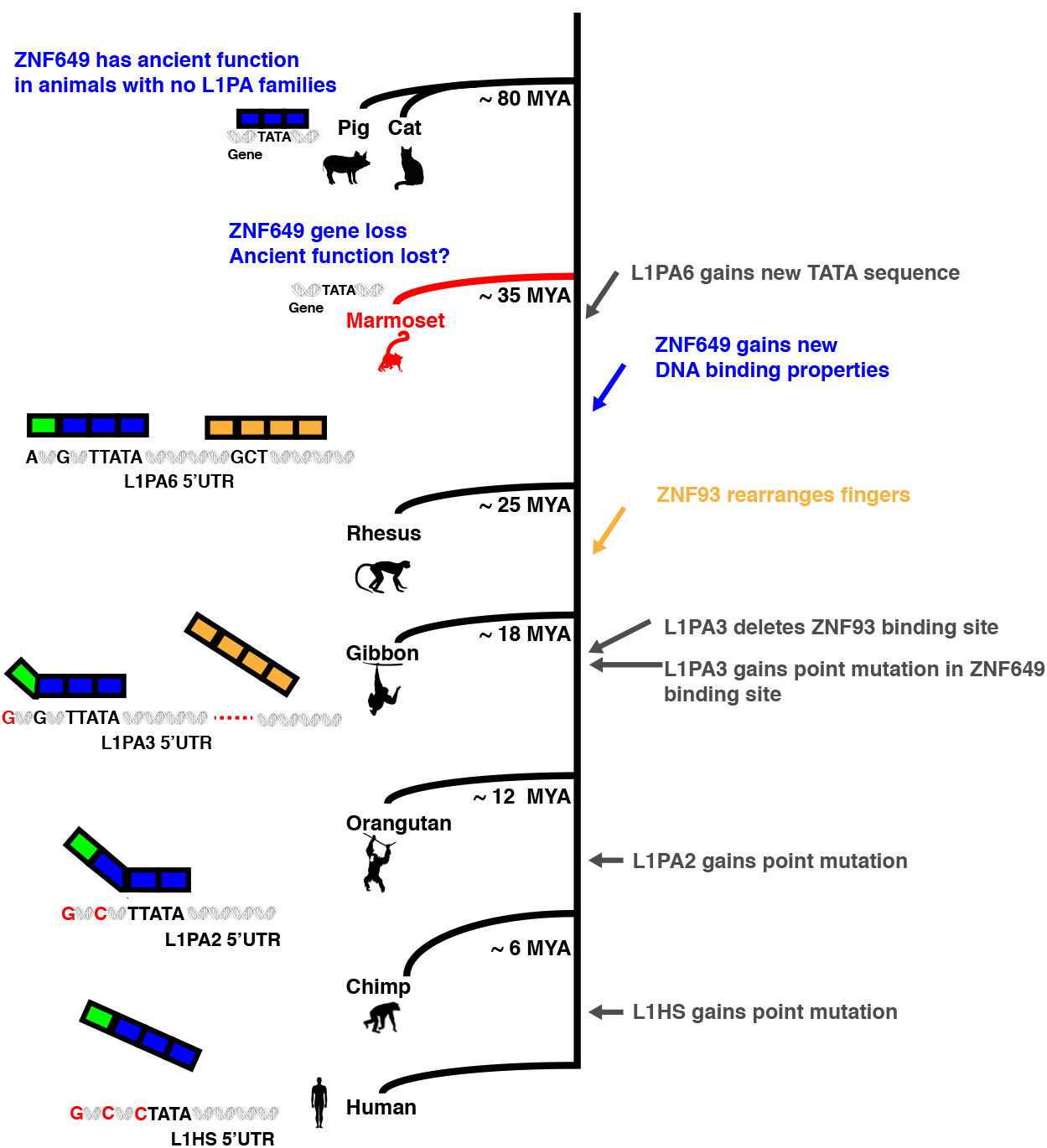
Model for L1PA arms races with ZNF649 and ZNF93. (Top) ZNF649 (blue) is found in cats and pigs where it presumably regulates cellular genes as these animals have no L1PA6 elements. Marmosets (red) lose ZNF649, and presumably any ancient ZNF649 function. After the marmoset divergence, L1PA6 elements invade the primate lineage leading to ZNF649 and ZNF93 evolution that results in both KZNFs binding and repressing the RTE together. L1PA3 elements subsequently acquire the deletion of the entire ZNF93 binding site and a point mutation that loosens the ZNF649 binding site and is eventually followed by two successive mutations in L1PA2 and L1HS elements that allow complete escape from ZNF649 binding.

These results illustrate novel mechanisms of evolutionary innovation, whereby a host genome rapidly evolves unique “teams” of KZNFs with distinct DNA binding abilities in order to repress RTEs. Some team members such as ZNF649 target essential portions of the RTE, with RTE escape requiring gradual point mutational paths on which a ZNF’s grip slowly “loosens” finger by finger; other team members, such as ZNF93, play supporting roles and target non-essential portions of the RTE that are rapidly escaped via a single event in which the ZNF completely “drops” the TE. Furthermore, ZNF649 may have been itself repurposed to battle L1PA, as previous literature demonstrates that it binds and regulates genes in highly conserved and cellular growth factor pathways^16^, possibly explaining its role in species that never faced L1PA6 elements. This repurposing demonstrates evolution’s exquisite ability to reuse existing cellular tools, and suggests that these genetic conflicts can compel ancient regulatory networks to evolve at the tempo of an arms race, which can then drive the creation of species-specific regulatory networks^17–19^.

## Methods

### UCSC Repeat Browser Analysis

We mapped Kap1 and KZNF ChIP-SEQ data to the UCSC Repeat Browser as previously described. To calculate the percentages of each element bound, we filtered all L 1PA7–2 and L1HS elements annotated in Repeat Masker to only full length elements (size limits of 5500–7500 nt) and used bedtools to intersect these instances with ChIP-SEQ datasets.

### Determination of Motifs

To perform discriminate analysis of motifs, we extracted 140 nt centered around the Repeat Browser meta-summit for every L1PA6, L1PA5, L1PA4, L1PA3, L1PA2 and L1HS element. We then used DREME^10^ to perform a discriminative analysis on these sequences using as our positive dataset all full-length genomic instances with ChIP-SEQ summits on them (bound), and the remaining instances as our negative set (unbound). We performed these comparisons for each L1PA family individually as well as a grouping of all L1PA6-HS elements together to confirm our predicted motif.

### mESC Reporter Assay

In order to generate luciferase constructs containing ZNF649 binding sites, we first synthesized a 201 bp region centered around our Repeat Browser metapeak on the L1PA4 consensus and cloned it into a Kpn I digested pgl-cp FFL vector (previously described^5^, map on Addgene) to create pglcp-SV40 ZNF649 L1PA4. We then used primers containing the appropriate point mutations to create pglcp-SV40 ZNF649 L1PA4–1mut (“L1PA3”), pglcp-SV40 ZNF649 L1PA4–2mut (“L 1PA2”), and pglcp-SV40 ZNF649 L 1PA4–3mut (“L1HS”). All maps will be provided on Addgene upon publication. To test the activity of each construct we plated E14 mESC at 200,000 cells/mL in a 24- well plate coated with 1% Porcine Gelatin. Cells were transfected 24 hours later with 100 ng of pCAG ZNF649 or pCAG Empty Vector with 20 ng of L1PA firefly luciferase reporter and 2 ng renilla luciferase. 24 hours later, cells were washed 2x in PBS and lysed for 15 min in 100 ul passive lysis buffer and 90 ul was analyzed per manufacturer’s instructions (Promega) on a Perkin Elmer 1420 Luminescence Counter.

### hiPSC Reporter Assay

To generate hiPSC knockdown lines for ZNF649, we designed guides downstream of the ZNF649 TSS using the CRISPOR track on the UCSC Genome Browser. Two separate guides with high efficacy and specificity were cloned into p783ZG, a modified version of MP783 (kind gift, S. Carpenter) in which the Puromycin-t2A-mCherry resistance gene was replaced by a Zeocin-t2a-GFP. Gen1C iPSC lines were then transfected with 1 ug of guide plasmid and selected at 50 ug/mL Zeocin for 2 weeks. The resulting stable populations were used with transient reporter. Briefly, two separate plates of the stable cell line pools were plated at 50,000 cells/cm^2^ and grown in 50 ug/mL zeocin (with one plate receiving 1 ug/mL dox). After two days dox-induced and uninduced cells were plated at 35,000 cells/well in separate Matrigel coated 24-well plates and transfected with 200 ng of the appropriate RTE reporter construct (as described above), 2 ng Nanoluc (Promega) and 2 ul Lipofectamine 2000 (Invitrogen). 24 hours later, cells were washed 2x in PBS and lysed for 15 min in 100 ul passive lysis buffer and analyzed per manufacturer’s instructions (Promega) on a Perkin Elmer 1420 Luminescence Counter.

### Generation and Testing of Alanine ZNF Mutants

We synthesized (Twist Biosciences) 10 codon optimized inserts (to break repetitive structure) containing an HA-tagged ZNF649 coding sequence, as well as the corresponding mutations to simultaneously ablate all DNA contact residues in a single finger in ZNF649. In addition to 10 independent single-finger mutants (1A, 2A, etc.), we synthesized a wild-type control. All constructs were tested on the “L1PA4” construct (ZNF649 binding site intact) in our mESC assay as described above.

### Reconstruction of ZNF649 Evolutionary History

The evolutionary history of ZNF649 was determined by syntenic and domain-based analyses as described in *Armstrong et al. in prep*. Briefly, we defined a high-quality syntenic locus encompassing ZNF649

## Acknowledgments

We thank all members of the Haussler lab for helpful conversations and constructive feedback, S. Carpenter and S. Covarrubias for CRISPRi plasmids and guidance in CRISPRi experiment design, B. Conklin for use of the Gen1C hiPSC line, and D. Kim for experimental suggestions. This work was supported by F32GM125388 to JDF. DH is an investigator of the Howard Hughes Medical Institute.

## Author Contributions

JDF, MH, and TT performed Repeat Browser analysis with assistance from SK. JDF, JA, BP, and DH analyzed the evolutionary history of ZNF649. JDF, KT, JG, NF, JP and PA performed experiments and analyzed data. JDF, SRS, and DH conceived of the project and designed the experiments and analysis.

## Author Information

No competing interests. Correspondence and request for material should be addressed to SRS and DH.

## Supplementary Information

**Extended Data Figure 1:**
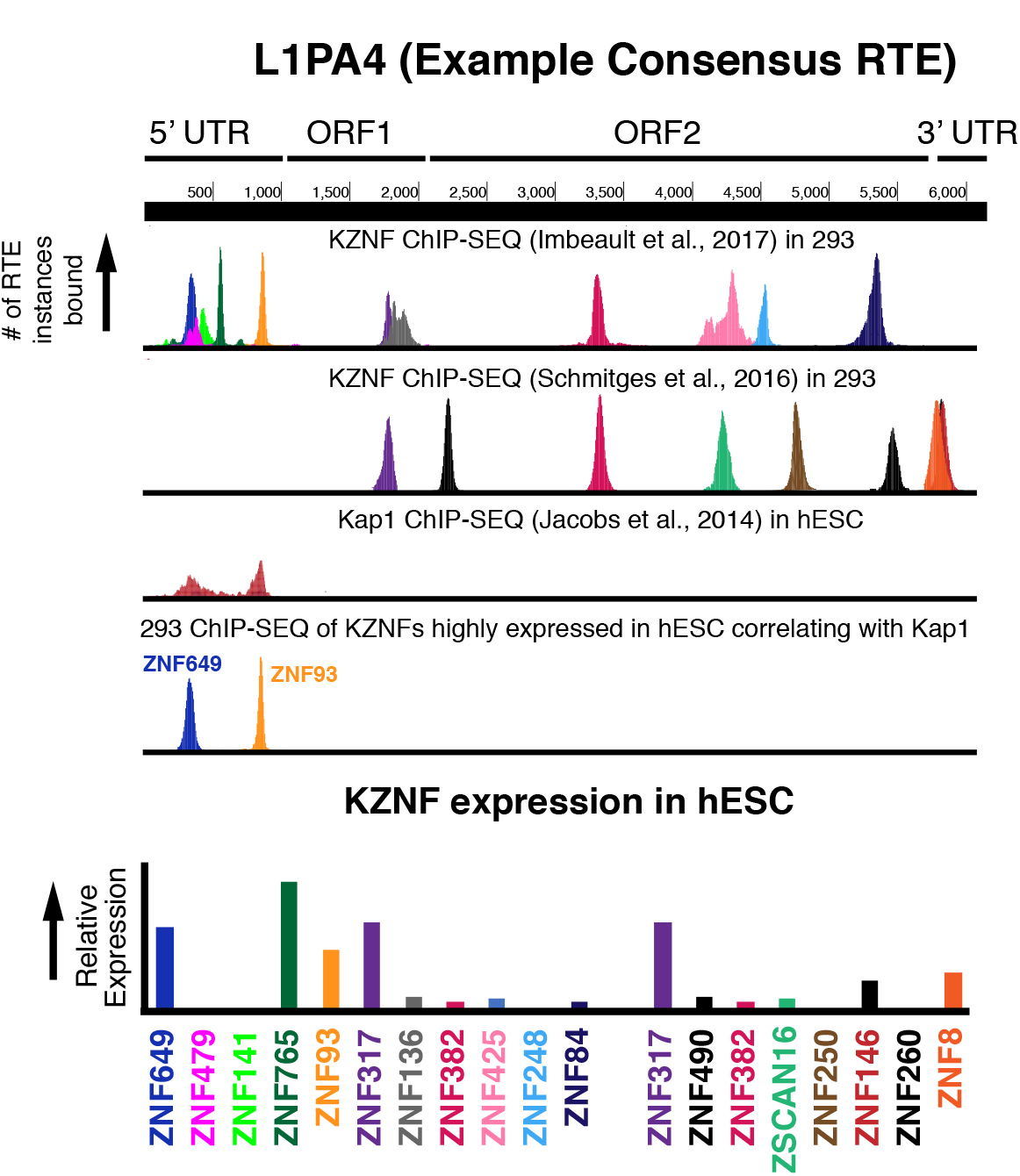
Mapping of all KZNF ChIP-SEQ to L1PA elements on the Repeat Browser (L1PA4 consensus shown as an example) shows many candidate KZNFs that might bind and repress these RTEs. However only ZNF649 and ZNF93 correlate with Kap1 binding and have high expression in pluripotent stem cells.

**Extended Data Figure 2:**
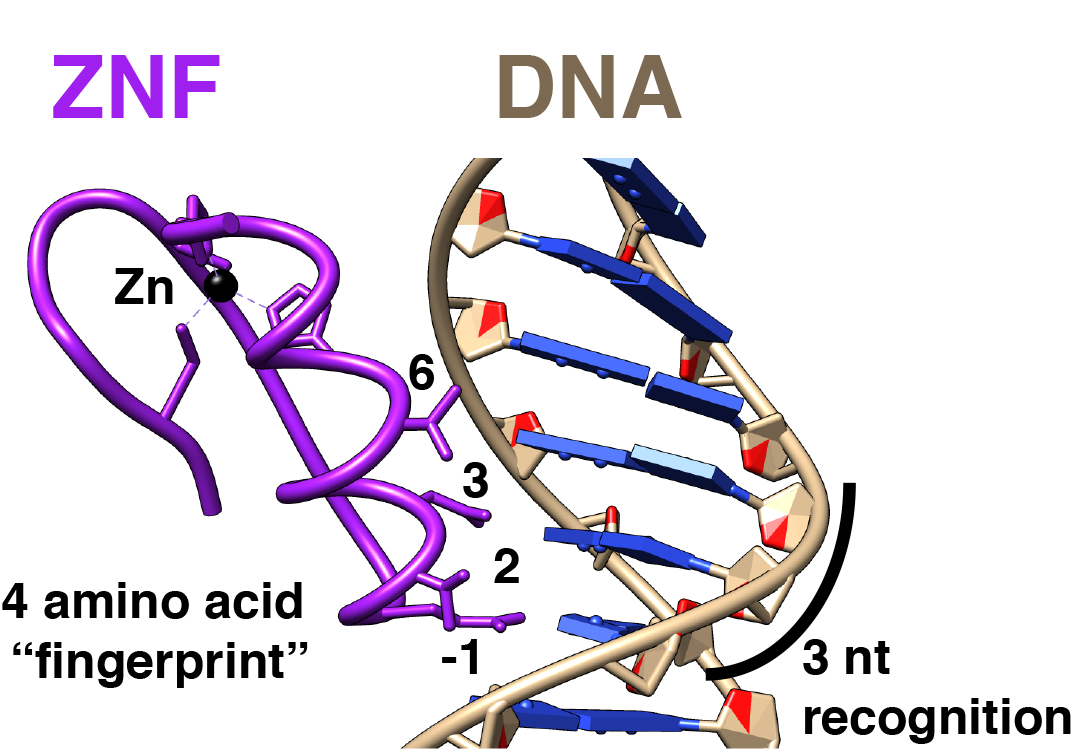
Canonical model for ZNF recognition on DNA. Shown here is the crystal structure (1G2F) of Zif268 bound to DNA. Numbered residues are in reference to the start of the helix (-1,2,3,6) and traditional make base-specific contacts to the DNA (four amino acid “fingerprint”).

**Extended Data Figure 3:**
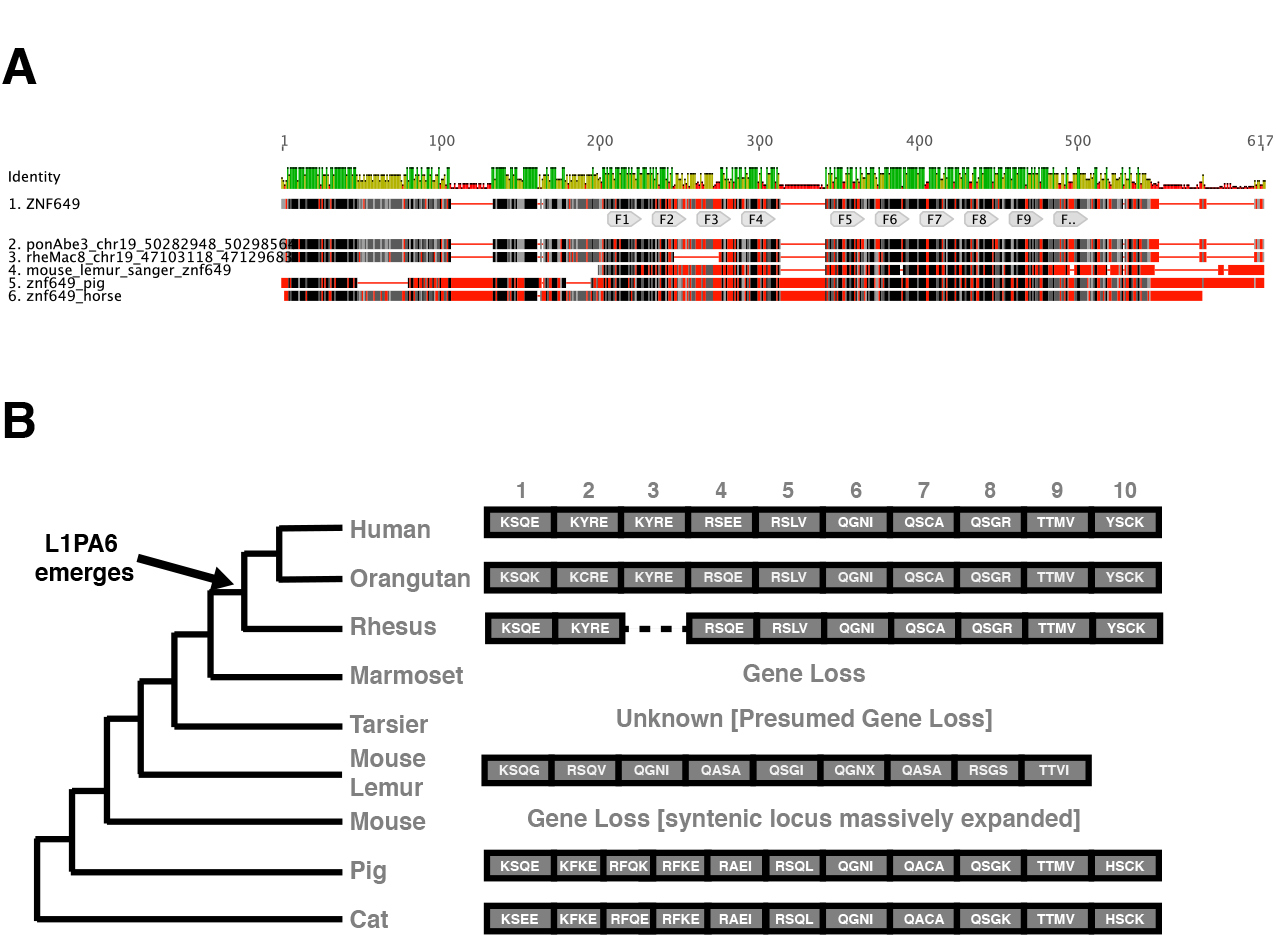
A) Alignment of ZNF orthologs as identified by syntenic analysis and resequencing of genomic DNA (mouse lemur). Fingers 1–10 (as defined in human ZNF649) are labeled on the alignment. B) Cartoon representation of ZNF orthologs reduced to the DNA binding residues of each ZNF (four amino acid “fingerprint”).

